# Tangled up in fibers: How a lytic polysaccharide monooxygenase binds its chitin substrate

**DOI:** 10.1101/2023.09.21.558757

**Authors:** Henrik Vinther Sørensen, Mateu Montserrat-Canals, Ayla Coder, Sylvain Prévost, Susan Krueger, Gustav Vaaje-Kolstad, Kaare Bjerregaard-Andersen, Reidar Lund, Ute Krengel

**Affiliations:** Department of Chemistry, University of Oslo, NO-0315 Oslo, Norway; Centre for Molecular Medicine Norway, University of Oslo, NO-0318 Oslo, Norway; Large-Scale Structures group, Institut Laue-Langevin, 71 avenue des Martyrs, 38042 Grenoble, France; Department of Materials Science and Engineering, University of Maryland, College Park, Maryland 20742, United States; Center for Neutron Research, National Institute of Standard and Technology, Gaithersburg, Maryland 20899, United States; Faculty of Chemistry, Biotechnology and Food Science, Norwegian University of Life Sciences (NMBU), NO-1340 Ås, Norway

**Author notes:** Correspondence: Henrik V. Sørensen; Ute Krengel. Henrik V. Sørensen, Department of Biomedical Science, Malmö University, SE-205 06, Sweden; Kaare Bjerregaard-Andersen, Ottilia vej 9, H. Lundbeck A/S, DK-2500 Valby, Denmark.

**Keywords:** bacterial adhesion, bacterial colonization, chitin complex, contrast matching, enzyme, GbpA, LPMO, secreted colonization factor, Small-Angle Neutron Scattering (SANS, bio-SANS), *Vibrio cholerae*

## Abstract

Lytic polysaccharide monooxygenases (LPMOs) are redox enzymes that bind to and oxidize insoluble carbohydrate substrates, such as chitin or cellulose. This class of enzymes has attracted considerable attention due to their ability to convert biomaterials of high abundance into oligosaccharides that can be useful for producing biofuels and bioplastics. However, processes at the interface between solution and insoluble substrates represent a major challenge to biochemical and structural characterization. Here, we investigated the four-domain LPMO from *Vibrio cholerae*, *N-*acetyl glucosamine binding protein A (GbpA), to elucidate how it docks onto its insoluble substrate with its two terminal domains. First, we developed a protocol that allowed GbpA and chitin to form a stable complex in suspension, overcoming incompatibilities of the two binding partners with respect to pH. Using contrast variation small-angle neutron scattering (SANS), after determining the neutron scattering contrast match point for chitin (47% D_2_O), we characterized the structure of GbpA in complex with chitin by SANS, and by electron microscopy. We found that GbpA binds rapidly to chitin, where it spreads out on the chitin fibers, and smoothens their surface. In some locations, GbpA binding induces the formation of protein-chitin clumps containing hundreds of GbpA molecules. Together, this suggests how the secretion of GbpA efficiently prepares the ground for microcolony formation by the bacteria.

## Introduction

Since the discovery of lytic polysaccharide monooxygenases (LPMOs) in 2010 (1), their ability to degrade crystalline cellulose, xylan or chitin has drawn massive interest for conversion of renewable biomaterials to biofuels. LPMOs are multi-domain proteins found in a wide range of organisms, including bacteria, fungi, algae and insects as well as viruses (2, 3). LPMOs are secreted by some human pathogenic bacteria and are employed for survival, both on carbohydrate surfaces outside and inside the host, using different strategies (4–6), including direct interference with the immune system (6). Given the abundance of these enzymes, and their sequence variation, it is likely that other important functions of these proteins are yet to be revealed. The discovery of LPMOs necessitated a reclassification of enzyme families in the Carbohydrate-Active enZYmes database (CAZY; http://www.cazy.org), where they are listed under ‘auxiliary activities’ families (AA9-11 and AA13-17). The first crystal structures of LPMO complexes with carbohydrate ligands (cellotriaose, Glc_3_ and cellohexaose, Glc_6_) were determined in 2016 by Frandsen *et al.* (7), for an AA9 LPMO from *Lentinus similis*. At approximately the same time, Courtade *et al.* probed ligand binding to another AA9 LPMO, from *Neurospora crassa*, by nuclear magnetic resonance (NMR) spectroscopy (8). These, and subsequent studies, showed how the ligand stretches across the flat surface of the pyramidal LPMO structure, with the +1 sugar unit binding to the N-terminal histidine of the characteristic copper-coordinating histidine-brace-motif present in the active site of LPMOs (Fig 1A). The catalytic mechanism of LPMOs, and current controversies, are summarized in an excellent review by Forsberg *et al.* (9). However, not all LPMOs can bind to oligosaccharides, and despite accumulating structural and functional data, there is scarce structural information on the molecular interaction of LPMOs with their complex, recalcitrant carbohydrate substrates. In particular, the complexes between the proteins and larger carbohydrate substrates such as chitin and cellulose have not yet been structurally characterized with experimental methods.

**Fig 1.**
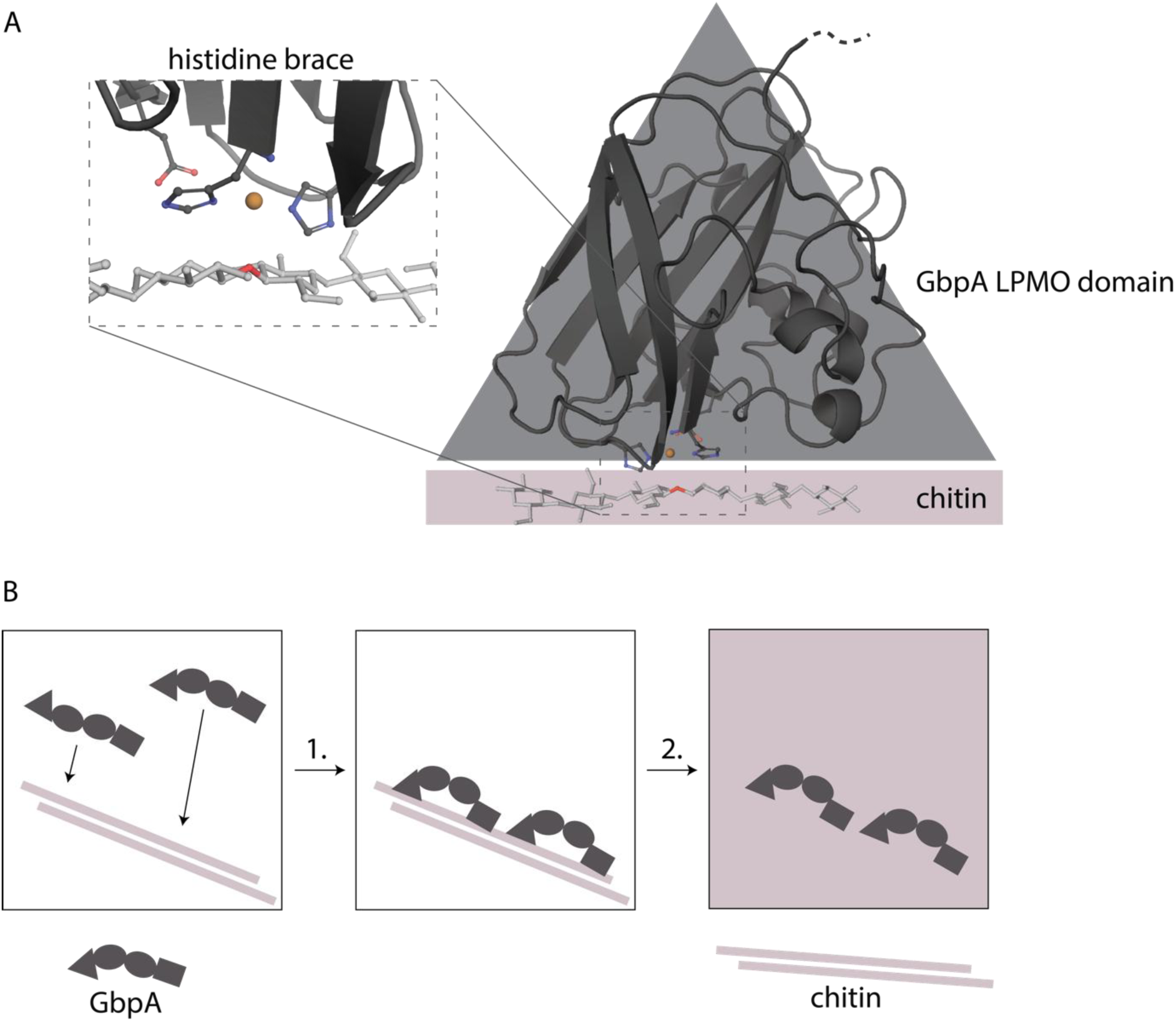
GbpA and schematic representation of SANS experiment. **A** The LPMO domain of GbpA binds to chitin and produces oxidative breaks in the fiber using its copper-containing histidine brace in the active site. Chitin has been manually modeled in its expected position based on (56), taking into account information from (10) (using PDB ID: 2XWX (15)). **B** Representation of SANS experiment with GbpA and chitin using contrast matching: (1) GbpA binds chitin, undergoing a conformational change; (2) by applying mixtures of D_2_O and H_2_O, chitin can be matched out in the SANS experiment, leaving GbpA as the only visible scatterer.

We are interested in virulence factors of *Vibrio cholerae,* the causative agent of cholera. Therefore, we selected *N-*acetylglucosamine binding protein A (GbpA) from *Vibrio cholerae* as a model system to study its interaction with chitin (for a review of chitin-active LPMOs, see Courtade & Aachmann (3)). GbpA was discovered in 2005 (4), and found to be important for colonization in the aquatic environment (4, 11, 12) as well as in the human host, binding to (and up-regulating) human mucins (5, 11). Recent studies further suggest that GbpA may interact with Toll-like receptors on host cells and induce an immune response (through its fourth domain) (13, 14), mediated by IL-8 secretion (13). GbpA consists of four domains, of which the first domain, which structurally resembles carbohydrate-binding module CBM33, is a chitin-degrading LPMO (15, 16) with calcium-binding properties (17), and additionally binds to mucins (15). The fourth domain (classified as a CBM73 in CAZy) also binds chitin (15), whereas domains 2 and 3, which structurally resemble flagellin and pilin-binding proteins, respectively, mediate attachment to the cell surface of the bacteria (15). The structure of the first three domains of GbpA has been solved by X-ray crystallography (15), recently complemented with the full-length crystal structure of a GbpA homolog from *Vibrio campbellii* (18). In addition, the solution-structure of GbpA has been characterized with Small-Angle X-ray Scattering (SAXS) and Small-Angle Neutron Scattering (SANS) (15, 19), revealing a monomeric elongated structure. None of the crystal structures contain carbohydrate ligands, and no structural data have been reported for the interaction between GbpA and chitin.

Here, we set out to reveal how GbpA interacts with chitin fibers using SANS. We characterized the bound and unbound GbpA structures by applying SANS to deuterated and non-deuterated GbpA (alone or in complex with chitin), and by varying the contrast using different D_2_O/H_2_O-mixtures, as schematically shown in Figure 1B. Additional, complementary insights were obtained from negative-stain electron microscopy (EM).

## Results

### Development of a protocol for solution studies of the GbpA-chitin complex

A suitable method for structural characterization of individual partners in a protein complex is SANS. This poses a challenge for studying protein complexes of chitin, which is an insoluble homopolymer of *N*-acetyl glucosamine (GlcNAc) residues. To our knowledge, chitin has rarely been used for solution scattering studies, because of its low solubility. It has, however, been shown that fine β-chitin nanofibers can be suspended in slightly acidic buffers (pH ∼ 3) (20). For shorter periods of time, chitin can be kept at pH ∼5, however, this is still low for GbpA, which shows some precipitation at pH ≤ 5. Fortunately, we could overcome these challenges by adding small volumes of 10-12 mg/mL GbpA (pH = 8.0) to chitin nanofibers (pH = 5.0), followed by immediate thorough pipetting and—for SANS experiments—sonication for 15-30 sec with a tip sonicator, being careful to avoid overheating, as described in the *Experimental procedures*. We observed that the mixture immediately became less viscous in this process. The samples treated in this manner were subsequently dialyzed into 20 mM sodium acetate-HAc pH = 5.0 during a 24-hour period. After this procedure, the chitin-GbpA complex could be kept in suspension for 12-24 hours, sufficient for several SANS experiments. For the lengthier USANS measurements, rotating tumbler cells were used to keep the samples in suspension. In the case of delays, the complex was re-sonicated shortly directly before the experiments. We observed that the solubility of the chitin fibers increased after the addition of GbpA, indicating that a complex had indeed formed. It was even possible to adjust the pH of the complex mixture to higher pH values (∼7), without immediate precipitation.

### Small-angle scattering of GbpA and chitin

Before exploring the structure of the chitin-GbpA complex, we investigated the structures of the two individual components. The solution structure of GbpA had already previously been characterized by SAXS and SANS (19), showing an elongated, flexible shape with a radius of gyration (*R_g_*) of 35-40 Å. Indeed, the scattering of GbpA cannot be described by a simple ellipsoid, but requires at least five connected ellipsoids to replicate the scattering data well (Fig 2A). Even though GbpA is a four-domain protein, this is not entirely surprising since the first domain is approximately twice the size of the other domains, and pyramidal in shape (not ellipsoidal).

**Fig 2.**
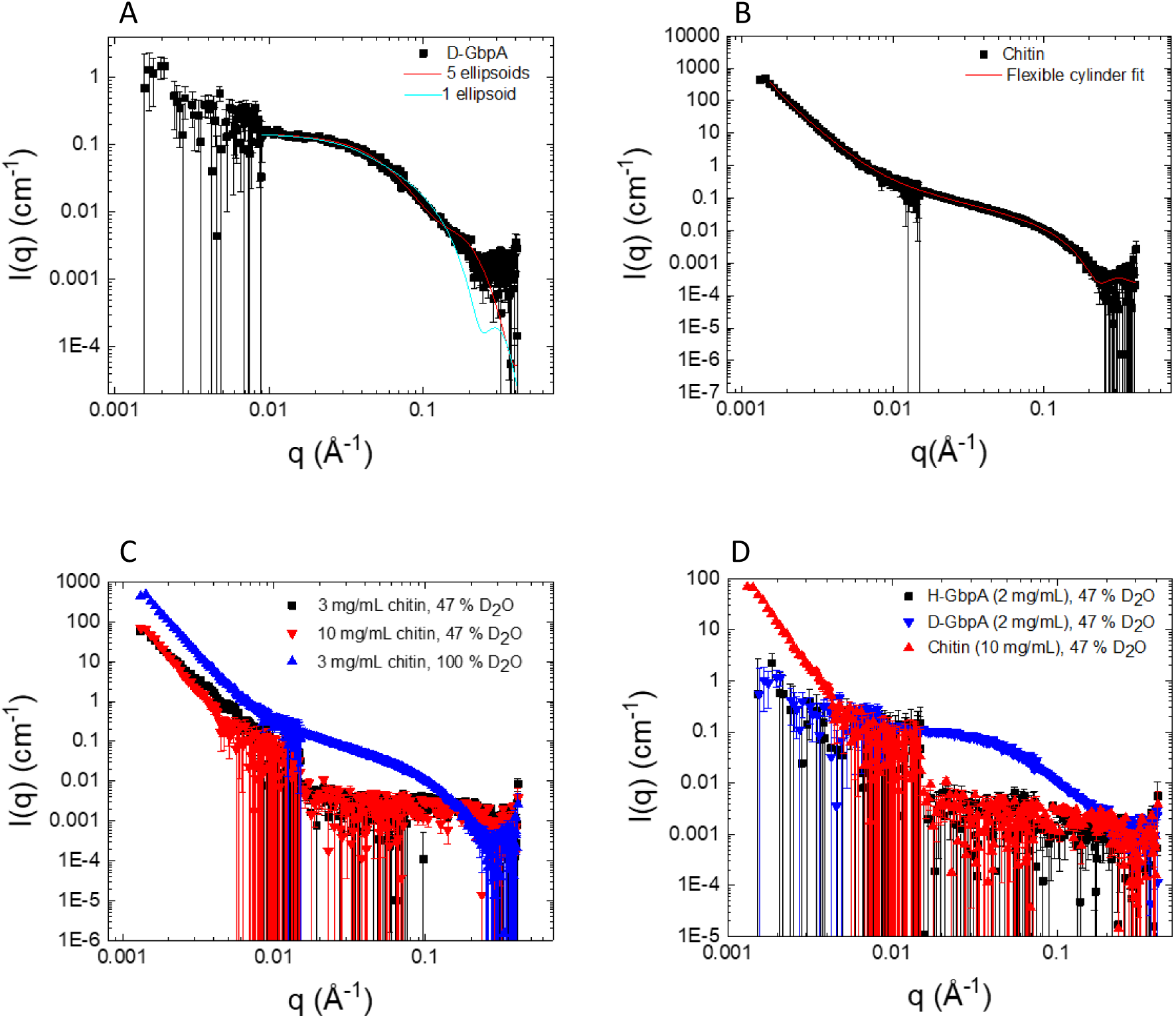
Independent scattering of GbpA and chitin. **A** SANS of D-GbpA in 47% D_2_O (the chitin match point). The protein is fairly extended in solution, hence describing the scattering as an ellipsoid (blue curve) gives a poor fit; a better fit can be achieved with 5 random-walk ellipsoids (red curve). **B** 3 mg/mL chitin was measured with SANS in D_2_O. The data can be fitted with a flexible cylinder model including a power-law for the steep upturn at low-q. **C** Scattering of chitin in 47% D_2_O is significantly weaker than in 100% D_2_O, and is matched out in the middle to high q-range. However, at low-q, chitin could not be matched out perfectly. **D** Comparing the scattering of D-GbpA in 47% D_2_O to chitin shows that the protein scatters very well in the middle to high q-range, where both chitin and H-GbpA are well matched out. Deuteration of the protein is thus essential to get sufficient contrast between the two components in an interaction study. Error bars represent one standard deviation.

Chitin fibers were modeled with a flexible cylinder model based on previous SAXS and SANS images of chitin fibers showing cylindrical shapes with some bends (21, 22). A similar approach has previously been used to model other nanofibers, such as proteins or cellulose; in the latter case, a slightly more complex variation was used, with an elliptical cross-section (23, 24). We found a circular cross-section to be adequate for chitin. To take into account chitin’s crystallinity and its sharp interfaces with the solvent, which give rise to a steep slope at low-q in SANS, we also included a power law in the model. The model fits the chitin data well (Fig 2B; χ^2^ = 2.78). The decay exponent in the power law approximates 4, which is consistent with sharp interfaces impenetrable for the solvent. Furthermore, the chitin fibers show a radius of 16.0 +/- 0.1 Å (Table S1), consistent with previous studies (20, 25). The model also gives the Kuhn length (318 Å; Table S1), which is twice the persistence length of the cylinder and thus relates to flexibility. A Kuhn length of 318 Å shows that chitin is very rigid. Both radius and Kuhn length are close to what has previously been reported for nanofibers of cellulose (23). The model can also yield the contour length (overall length of a fiber), but this parameter is too large to be meaningfully approximated from the q-range in SANS.

### Chitin can be matched out for SANS studies of GbpA-chitin complex

To obtain structural information of GbpA in its chitin-bound state with SANS, it is necessary to measure the sample at the chitin D_2_O/H_2_O match point. Online tools (26, 27) suggested a match point around 44% D_2_O for a chitin density of 1.5 g/ml, which is close to the theoretical match points of most globular proteins (∼40-45% D_2_O) and may thus potentially present a challenge for distinguishing chitin from interacting proteins. However, online tools only give estimates, and experimental determination is required. Moreover, chitin has a complex structure, where crystalline domains might have different densities than less crystalline regions and a small degree of deacetylation is expected. Additionally, residual components from the natural source may persist in some regions (divalent ions, fatty acids, *etc*. (28)), thus it was not even clear from the outset, if a defined match point could be obtained. We determined the chitin match point with SANS to be at 47% D_2_O (Fig S1). This necessitated the use of deuterated GbpA (D-GbpA) in order to distinguish the protein from chitin in the SANS interaction studies. To confirm that chitin is well matched out in the q-range where D-GbpA scatters, we measured chitin, D-GbpA and non-deuterated GbpA (H-GbpA) individually at 47% D_2_O (Fig 2C-D). The scattering of 2.5 mg/mL D-GbpA at 47% was much stronger than for chitin, which was well matched out, except at very low q-values.

### GbpA decorates chitin, thickening and smoothing the fibers

Following the establishment of the contrast match point, we progressed to study the complex of chitin and GbpA. An indication of complex formation was already obtained upon mixing the two components, as described above. The chitin suspension was highly viscous, but with the addition of D-GbpA and subsequent sonication, the suspension became easier to pipette and stuck less to surfaces (images of the chitin suspension and the mixture are shown in Fig 3A, B). Nevertheless, some sedimentation occurred, which affected reproducibility and prevented a more detailed analysis of the scattering data. The SANS curve of the complex at the chitin match point (Fig 3C, D) does not feature the characteristic GbpA form factor bump around q = 0.05 Å^-1^, confirming GpbA to be in the chitin-bound state, where an *R_g_* of 35-40 Å does not persist. Interestingly, the SANS curve has features similar to the SANS curve for chitin itself, with a good contrast (at 100% D_2_O). In particular, a steep slope at low q-values is present for both chitin and the complex mixture (Fig 3E), indicating that GbpA spreads out along the chitin fibers. The steep slope also indicates that the samples have a protein-protein structural correlation dictated by the chitin structure. These results were confirmed with negative-stain EM by analyzing GbpA and chitin mixed together. Here, the addition of full-length GbpA (GbpA_fl_) to chitin produces a change in the ultrastructure of the chitin fibers (Fig 4). The fibers appear less stiff and with less defined edges, indicating that GbpA coats the fibers.

**Fig 3.**
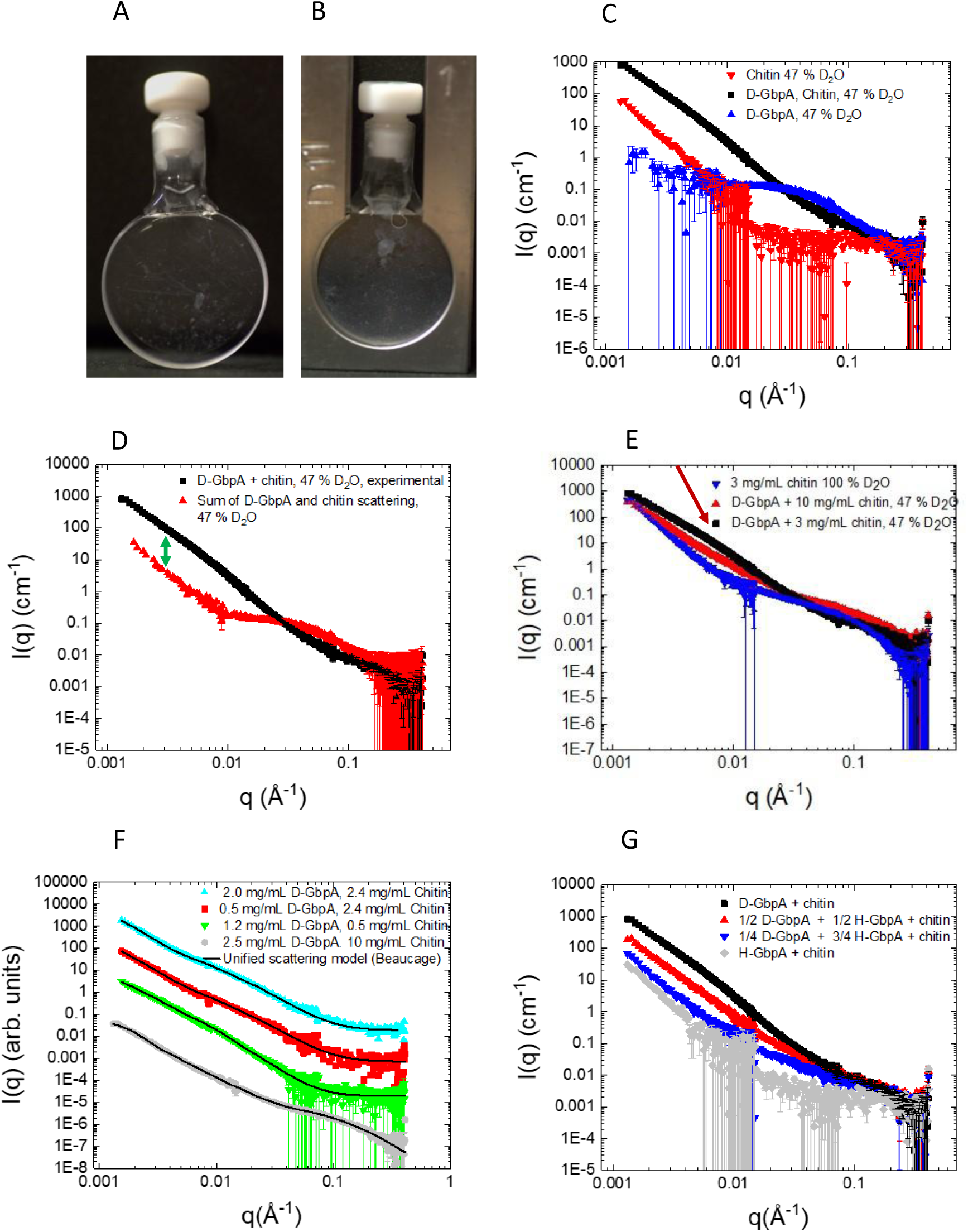
Sample preparation and SANS experiments. **A,B** Images of 3 mg/mL chitin samples, either without GbpA (A) or with 2.5 mg/mL GbpA (B). **C-G** SANS data of GbpA, chitin or GbpA-chitin mixtures, plotted as intensities (I(q)) vs. the scattering vector (q) on double-logarithmic scale. Unless stated otherwise, 3 mg/mL chitin (except samples of GbpA alone), 2.5 mg/mL GbpA (combined concentration of H-GbpA and D-GbpA; except for samples of chitin alone) was used and measurements were taken at 47% D_2_O. **C, D** Chitin and D-GbpA clearly form a complex, as the SANS data of the mixture strongly differ from the sum of the scattering of the individual biomolecules. The disappearance of the GbpA feature (large bump) around 0.05 Å^-1^ reveals a change in the GbpA structure upon chitin binding. The 20-30 fold increase in intensity at low q compared to the sum of GbpA and chitin scattering (arrow in D) suggests scattering by very large clusters of GbpA molecules, possibly GbpA bridging multiple chitin fibers together creating thicker layers. **E** Increasing the amount of chitin relative to GbpA changes the overall structure of the GbpA aggregate, but the steep slope at low-q persists, suggesting a structural correlation between the proteins. The bump at mid-q (indicated by red arrow), is related to changes in fiber thickness upon GbpA binding. **F** The SANS data of the GbpA-chitin complex can be fitted with a unified scattering model (see fitting parameters in Table 1 and discussion of the parameters in the main text). **G** Varying the H-GbpA to D-GbpA ratios affected the structure factor (intensities were lower for higher fractions of H-GbpA, since H-GbpA is almost matched at 47% D_2_O), but the steep slope at low-q persisted, showing that even at very low D-GbpA to H-GbpA ratios (*i.e.*, over long distances), the structural correlation between molecules does not disappear.

**Fig 4.**
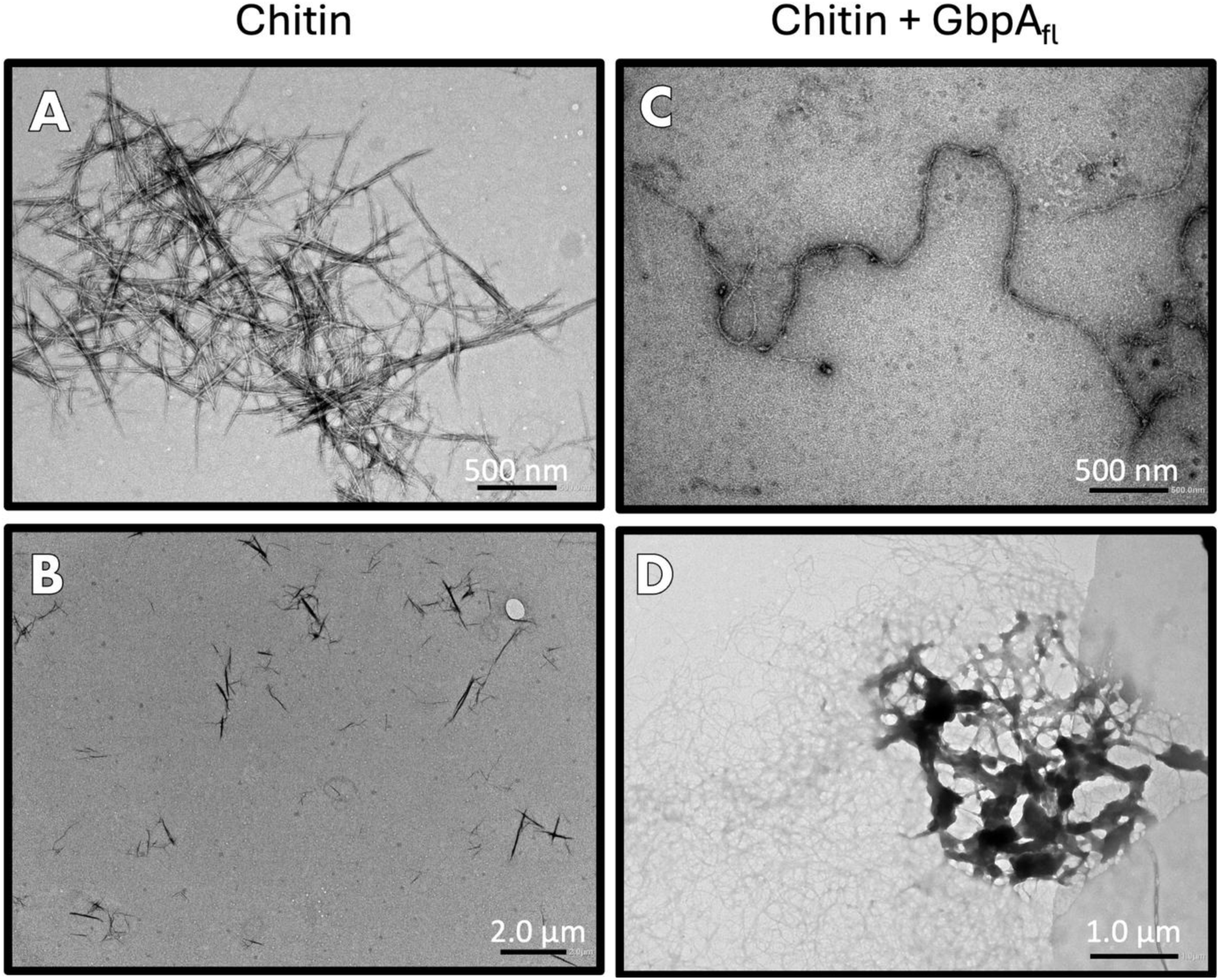
Negative-stain EM of chitin and GbpAfl. **A** Cluster of chitin fibers. **B** Overview image (zoomed out) of chitin fibers (uncoated). **C** Individual chitin fiber coated with GbpA_fl_. The structure is longer and smoother, like a string. **D** Representative dense aggregate of chitin fibers caused by GbpA_fl_. Individual fibers spread out from the cluster. The concentration of chitin in all conditions was 1.5 mg/mL, while the concentration of GbpA_fl_ (when present) was 400 µM. Extended TEM micrographs are shown in Fig S3.

**Table 1:** Fitting parameters, unified scattering model.

| Sample* | Decay<br>exponent 1 | $R_g$ 1 (Å) | Decay<br>exponent 2 | $R_g$ 2 (Å) | $\chi^2$ | SASBDB ID |
| --- | --- | --- | --- | --- | --- | --- |
| 2.5 mg/mL D-GbpA,<br>10 mg/mL chitin | $2.69 \pm 0.01$ | $1278 \pm 11$ | $3.88 \pm 1.1$ | $16.2 \pm 0.3$ | 2.66 | SASDW92 |
| 2.5 mg/mL D-GbpA,<br>3.0 mg/mL chitin | $2.64 \pm 0.01$ | $1859 \pm 71$ | $2.63 \pm 0.02$ | $125.2 \pm 0.2$ | 4.18 | SASDWA2 |
| 2 mg/mL D-GbpA,<br>2.4 mg/mL chitin | $3.64 \pm 0.05$ | $1600 \pm 169$ | $2.98 \pm 0.01$ | $202.5 \pm 0.3$ | 2.86 | SASDWC2 |
| 0.5 mg/mL D-GbpA,<br>2.4 mg/mL chitin | $3.40 \pm 0.17$ | $1633 \pm 387$ | $2.97 \pm 0.07$ | $183.1 \pm 12.0$ | 0.98 | SASDWD2 |
| 1.2 mg/mL D-GbpA,<br>0.48 mg/mL chitin | $3.16 \pm 0.14$ | $1484 \pm 210$ | $3.40 \pm 0.06$ | $221.8 \pm 6.9$ | 1.60 | SASDWF2 |
| 1.0 mg/mL D-GbpA,<br>1.2 mg/mL chitin,<br>72% D<sub>2</sub>O | $3.03 \pm 0.06$ | $1797 \pm 223$ | $2.36 \pm 0.60$ | $220.3 \pm 5.3$ | 1.03 | SASDWE2 |
| 1.25 mg/mL D-GbpA,<br>1.25 mg/mL H-GbpA,<br>3.0 mg/mL chitin | $2.81 \pm 0.01$ | $1845 \pm 131$ | $0.94 \pm 0.11$ | $43.4 \pm 1.8$ | 1.99 | SASDWH2 |
| 0.625 mg/mL D-GbpA, 1.875 mg/mL<br>H-GbpA, 3.0 mg/mL<br>chitin | $2.78 \pm 0.03$ | $2140 \pm 165$ | $0.43 \pm 0.11$ | $47.2 \pm 1.7$ | 1.95 | SASDWJ2 |

The data were fitted with Beaucage unified scattering model with two levels, yielding two $R_g$ s and two decay exponents. The first $R_g$ is not reliable, as the Guinier region is outside of the q-range, the first decay exponent relates to the fractal dimensionality of the aggregate. The second $R_g$ and decay exponent relates to structural features of the samples. An $R_g$ of 125-220 suggests several protein molecules contribute to the scattering. All datasets are available in the SASBDB, with the accession codes in the last column.
\*Samples are in 47% D<sub>2</sub>O unless stated otherwise.

Given the absence of clear features in the scattering that could be related to the protein form factor in the chitin-bound state, and the presence of a strong protein-protein structural correlation dictated by the complex chitin network, it was challenging to obtain a reliable structural model of the complex (a discrete structural model for similar systems does not exist to our knowledge). Initial attempts to model the data using the flexible cylinder model (as for chitin itself), did not yield satisfying fits (χ^2^ > 16). We further attempted to fit the data with a random-walk ellipsoidal model (21) to mimic the alignment of protein domains on the chitin fibers. This also did not fit the data well (χ^2^ > 20). However, we could fit the data using the unified scattering model by Beaucage (29) (χ^2^ between 0.43 and 3.88 for different experimental conditions; see Table 1), which does not yield a geometric model (cylinders, ellipsoids), but enables extraction of structural parameters such as *R_g_* values even for overlapping structural levels in hierarchical systems. The Beaucage model applies alternating power-law and Guinier regimes, yielding multiple radii of gyration for the different layers in the structure as well as insight into the structural nature of interfaces. This model has previously been used for several polymer systems (30). In some cases, this model can be used to obtain aggregation numbers, however this would require that one of the structural levels corresponds to a monomer and that the low-q Guinier region is covered in the q-range, which is not the case for our data. By analyzing the scattering data for the GbpA-chitin complex using the unified scattering model by Beaucage (Fig 3F, Table 1), we obtained *R_g_* values for the two structural levels within the samples. The larger *R_g_*-value (>1000) was generally unreliable, as the Guinier range was outside the limits, but the large size does confirm scattering from larger clusters. For most of the samples, an *R_g_* in the range 125-220 Å was obtained (compared to an *R_g_* ∼16 Å for undecorated chitin fibers; see Section *Scattering of GbpA and chitin*), likely related to the cross-section of the protein coating the fibers. Thickening of the fibers was also observed in transmission electron microscopy (TEM) images, obtained under similar conditions (Fig 4), although measurements of the fiber thickness in negative-stain EM are not very reliable due to variations in staining intensity between samples (even when the samples are stained for similar times).

### GbpA alters the chitin network structure and forms clusters

We observed that the overall scattered intensity (Fig 3D) increased about a 100-fold at low-q compared to the sum of chitin and GbpA scattering, indicating aggregation of GbpA on chitin at large length-scales. While the data do not contain a well-defined Guinier-region at low-q, a rough estimate can be made of the features around 0.01 Å^-1^ where the forward scattering (through extrapolation) is of the order of 200-2000 cm^-1^, which corresponds to a 100- to 1000-fold increase compared to the monomeric protein. This suggests that the scattering in the complex results from larger aggregates—comprising hundreds of protein molecules on the chitin network—rather than from GbpA monomers. Some aggregation can be induced by sonication, even using our careful approach (see *Experimental procedures*), however, the clusters were also prominent in the EM micrographs (visible around bundles of GbpA-covered chitin fibers; see Fig 4D), where the protein was never subject to sonication, suggesting chitin-induced GbpA clustering.

Interestingly, the *R_g_* is not completely reproducible, and a sample with lower protein:chitin ratio instead exhibits an *R_g_* of 16.0 Å, which is close to the radius of the chitin nanofiber chains alone (Table S1 and (20, 25)). This may be because GbpA largely spreads out on the chitin fibers unless GbpA is present in excess. However, similar results were not obtained at lower concentrations (while retaining the GbpA:chitin ratio), possibly due to a low signal at higher-q. The overall discrepancy in *R_g_* between samples can also be explained by different parts of the chitin network being in the neutron beam at the time of SANS data acquisition. SANS measurements of the D-GbpA–chitin complex at D_2_O levels between the match points of the two components (Fig S2) also yielded an *R_g_* of 220 Å for the protein aggregate; nevertheless, some of the characteristic features of chitin fibers are still present in the SANS data at high-q.

In addition to *R*_g_ values, the unified scattering model gives decay exponents for the different levels of structures (summarized in Table 1). The decay exponent describes the surface scattering or the fractal dimensionality of the aggregate (Porod-like) if D∼4 (31). Chitin itself has a decay exponent of 4.0 (Table S1), indicating very large aggregates with a sharply defined surface (31, 32) (as seen in Fig 4A). For the GbpA-chitin complex at the chitin match point, the low-q decay exponent is 2.5-3.6, consistent with aggregates in a state between random and globular. The smaller decay exponent compared to chitin suggests a partial loss of the sharp interfaces of the chitin fibers when decorated with protein, as seen by EM (Fig 4C). The decay exponent corresponding to the structural levels with *R*_g_ values of 125-220 Å is approximately 2.6-3.2, and thus considerably higher than 2.0 (which would be expected for an ideal random-walk chain). This shows that the protein-coating is rigid and compact.

### Individual GbpA molecules bound to chitin could not be resolved

By decreasing the ratio of GbpA to chitin, we attempted to spread out the GbpA molecules in order to analyze the structure of individual GbpA molecules bound to chitin. This did not work. To address this problem in a different way, we used mixtures of H-GbpA and D-GbpA bound to chitin at the chitin match-point of 47% D_2_O (Fig 3G). Since H-GbpA is well matched-out at the chitin match-point, we expected to still mostly see D-GbpA on the chitin fibers. However, since D-GbpA and H-GbpA compete for the binding sites, the distance between D-GbpA on the fibers should increase, perhaps to the point of scattering independently. As expected, the SANS intensities for the mixtures of D-GbpA and H-GbpA on chitin decreased with a higher proportion of H-GbpA (Fig 3G). Using the unified scattering model, an *R_g_* of 43-47 Å was obtained, which is close to that of monomeric GbpA; however, no clear features could be observed in the scattering curve in the corresponding q-range (∼0.05 Å^-1^) and overall, the data quality at high-q was significantly weaker for these samples due to the lower signal-to-noise ratio. While the low *R_g_* likely originates from the GbpA-layer on chitin, the actual value may not be indicative of the thickness, as some of it corresponds to matched-out H-GbpA. The decay exponent at low-q was 2.8, similar to the other GbpA-chitin samples; thus this approach did not remove the structure factor effect. This finding showed that even when the distance between D-GbpA particles on chitin is increased, D-GbpA molecules do not scatter independently (further increasing the distance between D-GbpA molecules by lowering the D-GbpA to H-GbpA ratio was not possible, since the signal became too low). Due to the rigid structure of the chitin network, it is likely that protein-protein structure factor effects will exist even at extremely low protein concentrations, making it technically impossible to determine the conformation of GbpA on chitin with SANS.

### GbpA binds to large areas of chitin rapidly and remains firmly attached

To further characterize the GbpA aggregates on chitin, we performed Ultra-Small-Angle Neutron Scattering (USANS) experiments, which give structural information in the hundreds of nanometer range, thus providing an estimate of the size of aggregates. However, we observed that the slope for the chitin-bound GbpA extends to very low angles without flattening out, indicating that GbpA decorates very large areas (or domains) on the chitin fibers (beyond the resolution of several micrometers; Fig 5). EM analysis provided a good explanation for this, with GbpA completely coating chitin fibers and inducing aggregation of the fibers in clumps as observed in Fig 4D. A more complete set of EM micrographs can be found in Figs S3 and S4. Interestingly, when experimenting with different GbpA:chitin ratios, we found that it made a difference if we prepared samples with individual ratios independently or if we started out with a 1:1 mixture to which we subsequently added either chitin or GbpA (Fig S5). In the latter case, we saw uncomplexed, rigid chitin fibers next to smooth GbpA-decorated chitin fibers with dense protein aggregates when adding chitin to the 1:1 mixture, suggesting that GbpA binds to chitin rapidly, covering its surface until all binding sites are occupied, and remains firmly attached to chitin after this.

**Fig 5.**
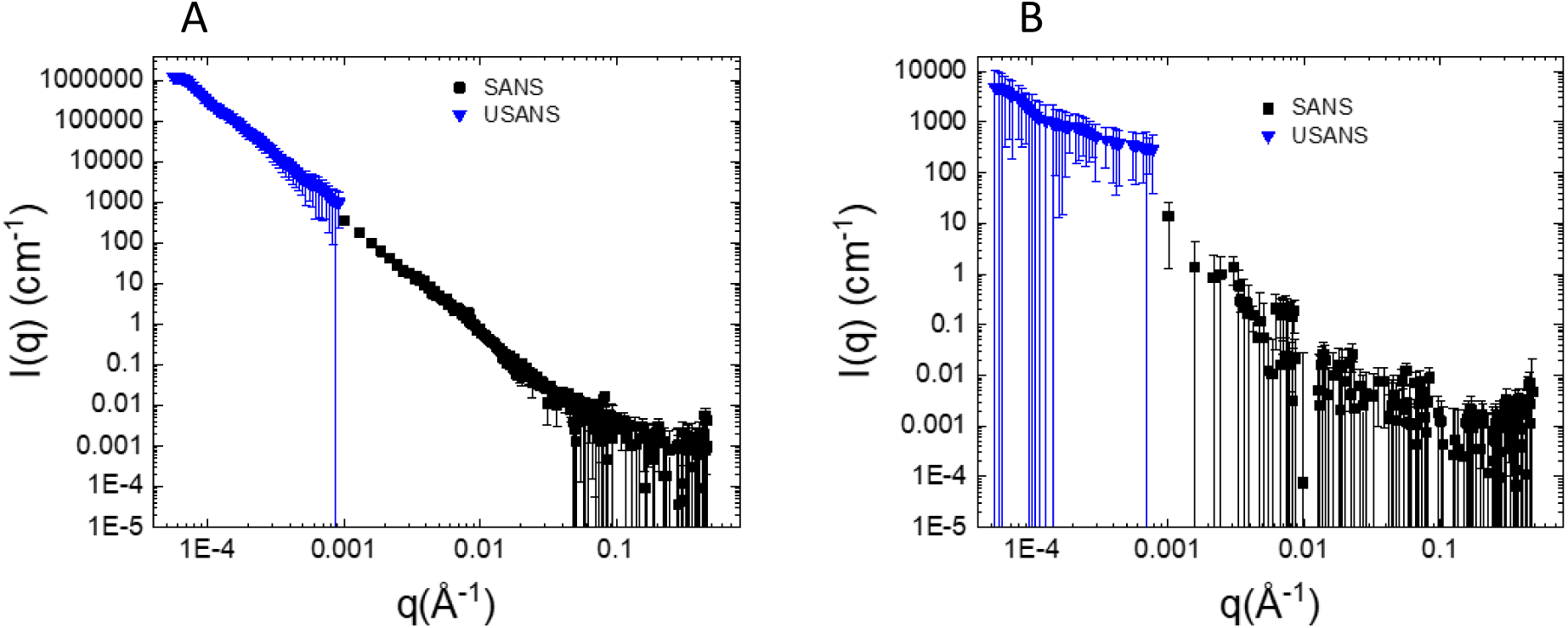
SANS and USANS data. **A** GbpA bound to chitin and **B** Chitin alone. Data were collected at 47% D_2_O. USANS data confirmed a structural order in the micrometer regime. Error bars represent one standard deviation.

### EM analysis of truncated GbpA and *V. cholerae* microcolonies

We were curious about the contributions of the different GbpA domains to chitin binding and complex formation, and therefore extended the EM analysis to truncated constructs of GbpA. Since both terminal domains (D1 and D4) are known to bind chitin (15), we selected constructs consisting of domains 1 to 3 (GbpA_D1-3_) and domain 1 alone (GbpA_D1_) for comparison with full-length GbpA (GbpA_fl_). GbpA_D1-3_ induced chitin aggregates similar to GbpA_fl_ (Fig 6B compared to Fig 4D). However, GbpA_D1_ appears to have a weaker effect on the fibers, as seen in Fig 6C and D. While the fibers are still decorated with the protein, they have much sharper edges compared to GbpA_D1-3_ and GbpA_fl_, and are visually more similar to the free chitin ultrastructure (Fig 4A and Fig S3), suggesting that domains 2-3 contribute importantly to GbpA network formation on chitin. A more complete set of EM micrographs can be found in Figs S6 and S7.

**Fig 6.**
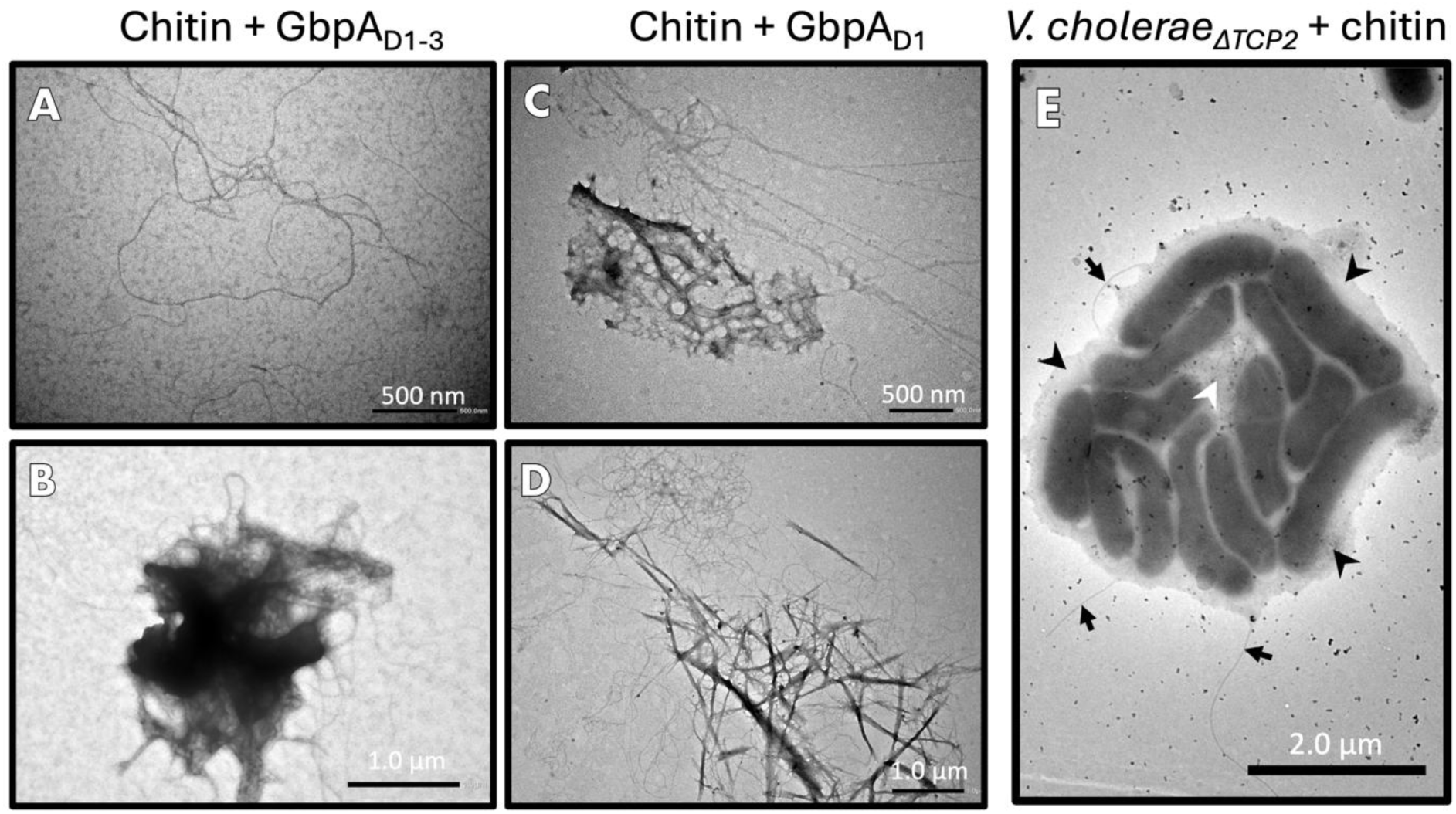
Extended negative-stain EM experiments for GbpA truncation variants and *V. cholerae* cells. **A** Individual chitin fibers coated with GbpA_D1-3_ truncation constructs. As for full-length GbpA, the fibers have a smooth appearance. **B** Cluster of aggregated GbpA_D1-3_-coated chitin fibers. **C** Individual GbpA_D1_-coated chitin fibers extend from an aggregation of fibers and protein. **D** Cluster of chitin bound to GbpA_D1_. The concentration of chitin in the experiments shown in panel A-D was 1.5 mg/mL, while the concentration of GbpA_D1-3_ and GbpA_D1_ in the experiments shown in A & B and C & D, respectively, were 400 µM (both cases). **E** Microcolony of *V. cholerae* observed only for bacteria incubated with chitin. A matrix of extracellular polysaccharides is indicated with black arrowheads; the hazy, fibrillar mass in the center, likely formed by chitin fibers, is indicated with a white arrowhead. Flagella are labeled with smaller black arrows. The concentration of chitin in this experiment was 1.5 mg/mL, while the bacterial culture had an OD_600_ of 2.0.

Additional EM experiments were performed by adding bacteria to chitin fibers, using an inactivated strain of *V. cholerae*, in which the toxin co-regulated pilus (TCP) was deleted. Interestingly, some of the bacteria were found to create microcolonies (Fig 6E) that were not observed for the bacteria imaged alone.

## Discussion

We developed a suspension protocol for the GbpA-chitin complex and carried out SANS, USANS and negative-stain EM experiments to characterize the structure of GbpA upon complex formation, allowing the first structural investigation of GbpA in the chitin-bound state. We observed that upon GbpA binding to chitin, the suspension of chitin-fibers immediately becomes less viscous, suggesting that the properties of the GbpA-chitin complex differ significantly from those of both partners alone. This was confirmed by SANS (Fig 3). Using EM, we observed that GbpA binding to chitin smoothed the otherwise rigid chitin fibers (Fig 4). After binding, GbpA remained firmly attached, and subsequent serial dilution by either chitin or GbpA had little effect on the formed complex (Fig S5).

In the SANS and USANS experiments, we experienced limits regarding the structural modeling. We suspect that it may not be possible to obtain a detailed structural model of independently scattering GbpA particles on β-chitin fibers using SANS, due to persisting structural order along the chitin fibers. This is not surprising, since the fibers are long and likely rigid, inducing a fixed order even over long distances. Much shorter chitin fibers would be required in order to overcome these technical impediments, if possible at all.

Both SANS and EM revealed that GbpA does not bind evenly to all parts of the chitin network (Fig 3 and 4), possibly because some parts of the fibers are less accessible than others. The regions of chitin with high accessibility for GbpA-binding likely exhibit protein-protein interactions between GbpA molecules yet to be elucidated. This hypothesis is supported by the weakened effect that shortened forms of GbpA (*i.e.*, GbpA_D1_) have on the chitin ultrastructure (as shown in Fig 6A-D). For example, domain 1 alone does not appear to be able to induce the formation of larger chitin aggregates (Fig 6C, D). This lessened interaction cannot be fully explained by the loss of the chitin-binding fourth domain, however, as the qualitative, *visual* effect of GbpA_D1-3_ on chitin was similar to GbpA_fl_ (Fig 6B vs. Fig 4D). Domains 2 and 3 therefore appear to be highly important for the observed clustering effect. These domains have previously been reported to be important for binding to the bacteria (15), thus they may have a dual role.

### Implications for GbpA’s role as colonization factor

Chitin is the most abundant biopolymer in marine environments, and is found, *e.g.*, on zooplankton, mussels and crustaceans. *V. cholerae* would therefore be well served by using a chitin-binding colonization factor, *i.e.*, GbpA, as one of its first anchors when forming microcolonies (as suggested by (33)). Other important factors for microcolony formation are the toxin-co-regulated pilus (TCP) (34, 35), the chitin-regulated pilus (ChiRP) (36) and outer membrane adhesion factor multivalent adhesion molecule 7 (Mam7) (37). An extracellular matrix consisting of polysaccharides and proteins then encases the microcolonies, completing the protected environment by forming a biofilm (38–41), which can adapt to changing environmental conditions (42, 43) and raises infectivity (44). Such a complex may be influenced by calcium, often found in mineralized chitin (17, 42). A negative-stain TEM image of a microcolony from an inactivated, TCP-deleted strain of *V. cholerae* is presented in Fig 6E (revealing that TCP is not essential for microcolony formation). Microcolonies were not observed in the absence of chitin. Chitin binding may thus initiate a cascade of events that culminate in biofilm formation and hyper-infectivity. Moreover, chitin binding has been shown to induce natural competence in *V. cholerae* (45), increasing bacterial fitness by exchange of genes. GbpA secretion appears to play an important role in the initial steps of these events. – But why would it make sense for the bacteria to throw out anchors without a connecting line and attach to them in a second step? The picture emerging from the present study is that GbpA efficiently prepares the ground for microcolony formation by forming aggregates dispersed throughout the fibers (Fig 7), and with this reserves space for *V. cholerae* over competitors, as previously suggested by Kirn *et al.* (4). Secreting anchors first, and then quickly attaching to them before reproducing, may allow the bacteria to very effectively colonize biotic surfaces in the aquatic environment. Additionally – and at least as important –, GbpA ensures access to food supplies through its lytic polysaccharide mono- (or per-) oxidase activity, giving the bacteria a solid return on their initial energy investment when producing and secreting GbpA.

**Fig 7.**
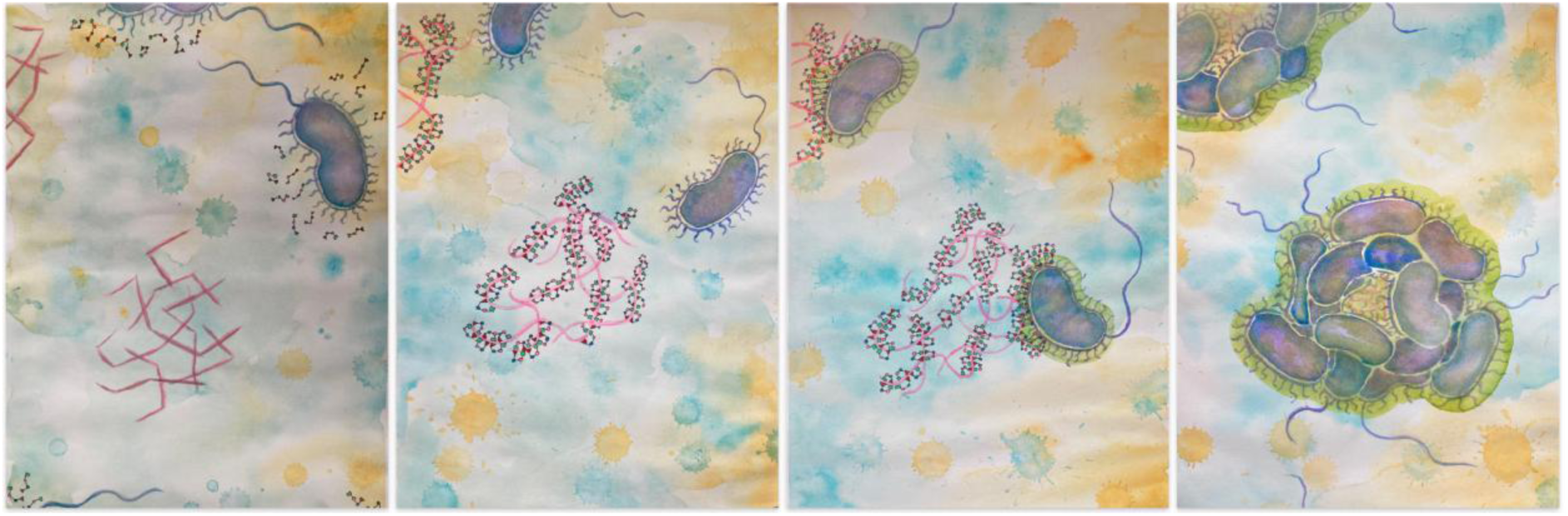
Water-color paintings of *V. cholerae* sketching microcolony formation on chitin aided by GbpA. *From left to right*: 1. *V. cholerae* bacterium secretes the four-domain protein GbpA (domains shown in different colors) close to a cluster of chitin fibers (pink, ragged appearance); 2. GbpA decorates chitin by attaching with its two terminal domains (LPMO-domain 1 (red triangle) and domain 4 (green)), smoothing the fibers’ appearance; 3. Bacteria attach via GbpA domains 2 and 3 (blue, pink) to GbpA-coated chitin fibers and produce a hazy, fibrillar mass (green halo around the bacteria), initiating biofilm formation; 4. Formation of *V. cholerae* microcolony attached to chitin via GbpA. (Credit: Ayla Coder)

### Experimental procedures

#### Protein production and deuteration

Perdeuteration of GbpA involved expression in deuterated media, according to the protocol described in (19), which was inspired by (46). This protocol relies on BL21 star (DE3) cells containing the GbpA-encoding gene in the pET22b vector (19). Growth and expression media contained D_2_O and d_8_-glycerol (99% D) purchased from ChemSupport AS (Hommelvik, Norway). The production of H-GbpA and the purification of D-GbpA and H-GbpA is also described in (19). The constructs of the truncated variants of GbpA (GbpA_D1_ and GbpA_D1-3_) for EM analysis are described in (15, 19), and the protocol for protein production and purification is almost the same as that used for the full-length construct, described in (19). The only difference concerns the purification of GbpA_D1-3_, where a step gradient was introduced during anion-exchange chromatography to improve the yields of pure protein. Specifically, a step corresponding to 180 mM NaCl (8 column volumes) was added to the isocratic gradient elution.

GbpA was saturated with copper to activate the enzyme prior to the SANS experiments, by incubation with CuCl_2_ in 5-fold molar excess for at least 30 min. Free copper was then removed by passing the protein through a HiTrap Desalting column (GE Healthcare). Prior to the SANS experiments, the protein was dialyzed against 100 mM NaCl, 20 mM Tris-HCl pH 8.0 with 47% of the water being D_2_O.

#### Preparation of chitin, GbpA and chitin-GbpA samples for SANS

β-chitin nanofibers (∼180 micrometer in length) from squid pens (France Chitine, Orange, France) were kindly provided by Jennifer Loose and Gustav Vaaje-Kolstad and prepared according to the protocol described in (16), which was based on the original study by Fan *et al.* (20). The fibers were dissolved in 20 mM acetic acid (pH 3.2) to a concentration of 10 mg/mL and subsequently sonicated at 50-70% power for 10-15 min (3 seconds on, 3 seconds off) with a Vibra Cell Ultrasonic Processor until they were well dispersed for several days. Finally, the fiber suspension was dialyzed against 20 mM sodium acetate-HAc pH 5.0 with the desired amount of D_2_O for at least 24 hours. At higher pH or in the presence of salts, it was observed that the fibers precipitated. However, at pH 5, GbpA is not very stable. Nevertheless, it was possible to add GbpA to the fibers from a highly concentrated stock solution of the protein (prepared by dialysis against 10 mM NaCl, 20 mM Tris-HCl pH 8.0), if the mixture was immediately treated by thorough pipetting (*i.e.*, pipetting the suspension up and down 7-10 times with a micro pipette, using a cut-off 1 mL pipette tip). To further stabilize the suspension, we sonicated the mixture for a total of 15-30 sec, 1 sec on, 1 sec off with a tip sonicator at 30-50 watts, avoiding overheating (for most samples, 15 sec total was sufficient), and re-dialyzed the sample for 24 hours against 20 mM sodium acetate-HAc pH 5.0 with relevant amounts of D_2_O. After any dilution of the chitin fibers or the mixture, the solutions were re-sonicated for 15-30 seconds to ensure proper suspension, but avoiding a raise in temperature of the sample. For GbpA-chitin mixtures, this step was only relevant for the measurement at 72% D_2_O. In case of delays before measurements, samples were re-sonicated for 2 sec.

GbpA samples without chitin were dialyzed into 100 mM NaCl, 20 mM Tris-HCl pH 8.0, with the desired level of D_2_O prior to SANS analysis.

#### SANS measurements

Samples were measured with Small-Angle Neutron Scattering (SANS) at beamline D11 at Institut Laue-Langevin (ILL) (47), with wavelength λ = 5.6 Å and a wavelength spread of Δλ/λ = 9%. Data were acquired for the q-range of 0.0013 – 0.4102 Å^-1^. Ultra Small-Angle Neutron Scattering (USANS) experiments were carried out at beamline BT5 (48) at the National Institute of Standards and Technology (NIST) Center for Neutron Research, using λ = 2.4 Å and a wavelength spread of 6% Δλ/λ, for the q-range 0.00005 Å^-1^ to 0.001 Å^-1^. Tumbling cells were used for USANS measurements. To find the chitin match point, 10 mg/mL chitin was measured in 0%, 20%, 42%, 66%, 80% and 100% D_2_O for 15 min per sample at ILL beamline D11. The data were integrated from 0.03 to 0.4 Å^-1^, and the integrated data were fitted with a second-degree polynomial function. The chitin match point was estimated from the minimum of this function.

While SANS samples were measured for 2-3 hours, USANS measurements lasted 11 hours per sample. Buffer, empty cell and H_2_O measurements were used for background subtraction and calibration to absolute scale. Data were processed with software at the respective beamlines (49, 50). The scattering data of GbpA were fitted with a random-walk ellipsoidal model using five ellipsoids (see reference (21) and equations within). For chitin the scattering was fitted with a flexible-cylinder model as implemented in SASVIEW (51), including a power-law for fitting the contribution from smooth interfaces between chitin clusters and solvent.

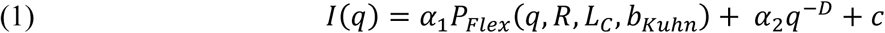

where α_1_and α_2_ are scaling factors, c is a constant background and D is the Porod decay exponent. *P*_*Flex*_ is the scattering from flexible polymers, depending on radius (R), contour length (*L*_*C*_) and Kuhn length (*b*_*Kuhn*_). *P*_*Flex*_ is modeled using the equations described by Pedersen and Schurtenberger (21), using their method 3 with excluded volume effects. We fixed the contour length to 5000 which is ∼2π/q_min_, to not model sizes outside the q-range. The data were uploaded to the Small Angle Scattering Biological Data Bank (SASBDB) (52), where they are accessible with accession code SASDW42.

For the complex of GbpA and chitin, the data were fitted with the Beaucage unified scattering model (29, 53):

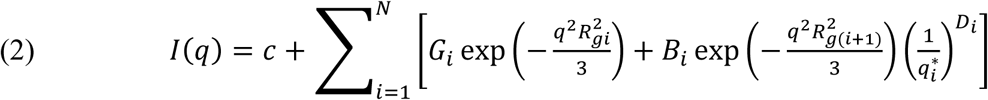

where *G* is the Guinier scaling factor, *Rg* the radius of gyration, *B* is the Porod constant and *D* the Decay exponent. *N* is the number of levels used and is set to two for all fits. *q*^*^_i_ is given by:

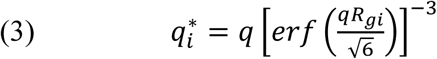

An upper limit of π/q_min_ was put on *R_g_* to keep values within the q-range. Two levels were first fitted individually to get starting parameters for the fitting of the levels combined. G and B are fitting parameters.

Data were plotted with Origin (Version 2017 64bit, OriginLab Corporation), and the modeling was carried out with SASVIEW 5.0.5 (51) and QtiSAS (54). The SANS data have been submitted to SASBDB (52), and the IDs are included in Table 1 and Table S1.

#### Negative-stain EM

GbpA and chitin samples were prepared, essentially as described above. 10 mL of 3 mg/mL chitin were subjected to sonication in 20 mM acetic acid at pH 3.2. Sonication was performed, on ice with a Q500 Sonicator (Qsonica), using the thinnest tip at 30% intensity in 3/3 sec on/off pulses until the sample was completely suspended. The solution was visibly inspected between rounds of sonication to ensure that the solution did not overheat, by only sonicating for 5-10 min at a time and waiting for 10 min between each sonication. Chitin was dialyzed overnight against 20 mM sodium acetate-HAc pH 5.0, and sonicated on ice again to ensure that it remained suspended. Chitin aliquots were then mixed with a few microliters of GbpA (or its truncated variants) at a final concentration of 400 µM, and incubated with agitation for 1h at 21 °C. Uncharged, carbon-coated 300 hex-mesh Cu TEM grids were exposed to 8 µL of sample, washed in deionized water 10x and stained with 1% uranyl acetate (UAc) for 30-60 seconds.

Additionally, two TEM experiments comparing different GbpA:chitin concentration ratios were performed. The chitin solution was re-sonicated to ensure full suspension for 10-15 minutes prior to use, using the same sonication method described above. These samples were mounted to 300 hex-mesh carbon-coated Cu TEM grids as described. In the first experiment (Fig S5A-C), two serial dilutions were carried out, starting from a 300 µL 1:1 solution of 1.5 mg/mL GbpA: 1.5 mg/mL chitin. In the first dilution series, a solution of 1.5 mg/mL GbpA_fl_ was used as the diluent to keep the concentration of GbpA_fl_ at 1.5 mg/mL while decreasing the chitin concentration. In the second dilution series, the diluent used was a 1.5 mg/mL chitin solution (to keep the chitin concentration at 1.5 mg/mL), while decreasing the GbpA_fl_ concentration. For both series, 30 µL of the 1:1 solution was first mixed by pipetting with 270 µL of diluent, before 30 µL from the resulting 1:10 (or 10:1) GbpA:chitin solution were immediately transferred to a new Eppendorf tube and mixed with 270 µL diluent. This was repeated two more times in each direction, creating the GbpA:chitin samples shown in Table S2. In the second experiment (Fig S5D-F), instead of performing a dilution series, the samples were prepared individually. Only three samples were prepared like this, as reported in Table S3.

Finally, TEM experiments were performed with an inactivated, TCP-deleted *V. cholerae* strain (*V. cholerae*_ΔTCP2_) and β-chitin. The bacterial samples were prepared by growing 100 mL liquid colonies in LB media overnight in a Multitron Standard shaker at 37 °C and 120 rpm. These bacterial samples (OD_600_ = 4) were mounted to 300 hex-mesh Cu TEM grids that were negatively charged in a glow-discharger. Each grid was applied to their respective sample drop (8 µL) for 30 minutes, then washed 5 times in distilled water and stained with 1.1% UAc for 30-60 seconds. All images were taken with a JEOL 1400Plus Transmission Electron Microscope operating at 120kV. Micrographs were analyzed with ImageJ (55).

## Data availability

SANS data are deposited in the Small Angle Scattering Biological Data Bank (SASBDB), with IDs given in Table 1. All other data are published in this manuscript.

## Acknowledgements

We thank Jennifer Loose for a very nice collaboration on the GbpA project and Hanne Winther-Larsen for providing the TCP-deleted *V. cholerae* strain. We further wish to thank the staff at NIST (Markus Bleuel, Paul Butler and Susana M. Teixeira) for outstanding beamline support and very fruitful discussions, and Mark Bleuel for preliminary comments on this manuscript. We are grateful to Jens Wohlmann and Norbert Roos for advice and support regarding EM grid preparation and image acquisition at the EM-lab at the Department of Biosciences, which is part of the UiO core facility *Life Science Electron Microscopy Consortium* (LSEMC). Other work at UiO was performed at the UiO Structural Biology core facilities, belonging to the Norwegian core facility NORCRYST.

## Author contributions

K.B.-A. and U.K. conceived the study. H.V.S. and M.M.-C. produced the protein, and H.V.S. established a protocol for sample preparation and performed SANS experiments, supervised by K.B.-A., R.L. and U.K., and with assistance of S.P. at ILL and S.K. at NIST. M.M.-C. and A.C. prepared the samples for EM and acquired the data at the EM core facilities at UiO. G.V.-K. supported the work with samples and know-how. The manuscript was written by H.V.S., with contributions from M.M.-C. and U.K.. It was revised with input and approval from all authors.

## Author contributions (CRediT Taxonomy)

Conceptualization: K. Bjerregaard-Andersen, U. Krengel

Data curation: H. V. Sørensen, M. Montserrat-Canals, Ayla Coder, S. Prévost, S. Krueger, R. Lund

Formal analysis: H. V. Sørensen, M. Montserrat-Canals, Ayla Coder, S. Prévost, S. Krueger, R. Lund

Funding acquisition: K. Bjerregaard-Andersen, U. Krengel

Investigation: H. V. Sørensen, M. Montserrat-Canals, Ayla Coder, S. Prévost, S. Krueger Methodology: H. V. Sørensen, S. Prévost, R. Lund

Project Administration: U. Krengel Resources: G. Vaaje-Kolstad Software: H. V. Sørensen, R. Lund

Supervision: K. Bjerregaard-Andersen, R. Lund, U. Krengel Validation: H. V. Sørensen, S. Prévost, R. Lund

Visualization: H. V. Sørensen, M. Montserrat-Canals, Ayla Coder Writing – original draft: H. V. Sørensen, U. Krengel

Writing – review & editing: H. V. Sørensen, M. Montserrat-Canals, Ayla Coder, S. Prévost, S. Krueger, G. Vaaje-Kolstad, R. Lund, K. Bjerregaard-Andersen, U. Krengel

## National Institute of Standards and Technology disclaimer

Certain commercial equipment, instruments, materials, suppliers, or software are identified in this paper to foster understanding. Such identification does not imply recommendation or endorsement by the National Institute of Standards and Technology (NIST) nor does it imply that the materials or equipment identified are necessarily the best available for the purpose.

## Abbreviations

CBM: carbohydrate-binding module
GbpA: *N-*acetylglucosamine binding protein A
D-GbpA: deuterated GbpA
EM: electron microscopy
H-GbpA: hydrogenated (= non-deuterated) GbpA
HAc: acetic acid
ILL: Institut Laue Langevin
LPMO: lytic polysaccharide monooxygenase
NIST: National Institute of Standards and Technology
PDB: Protein Data Bank
SANS: Small-Angle Neutron Scattering
SASBDB: Small Angle Scattering Biological Database
SAXS: Small-Angle X-ray Scattering
TEM: Transmission Electron Microscopy
USANS: Ultra-Small-Angle Neutron Scattering
UAc: uranyl acetate.

## SUPPORTING INFORMATION

**Fig S1.**
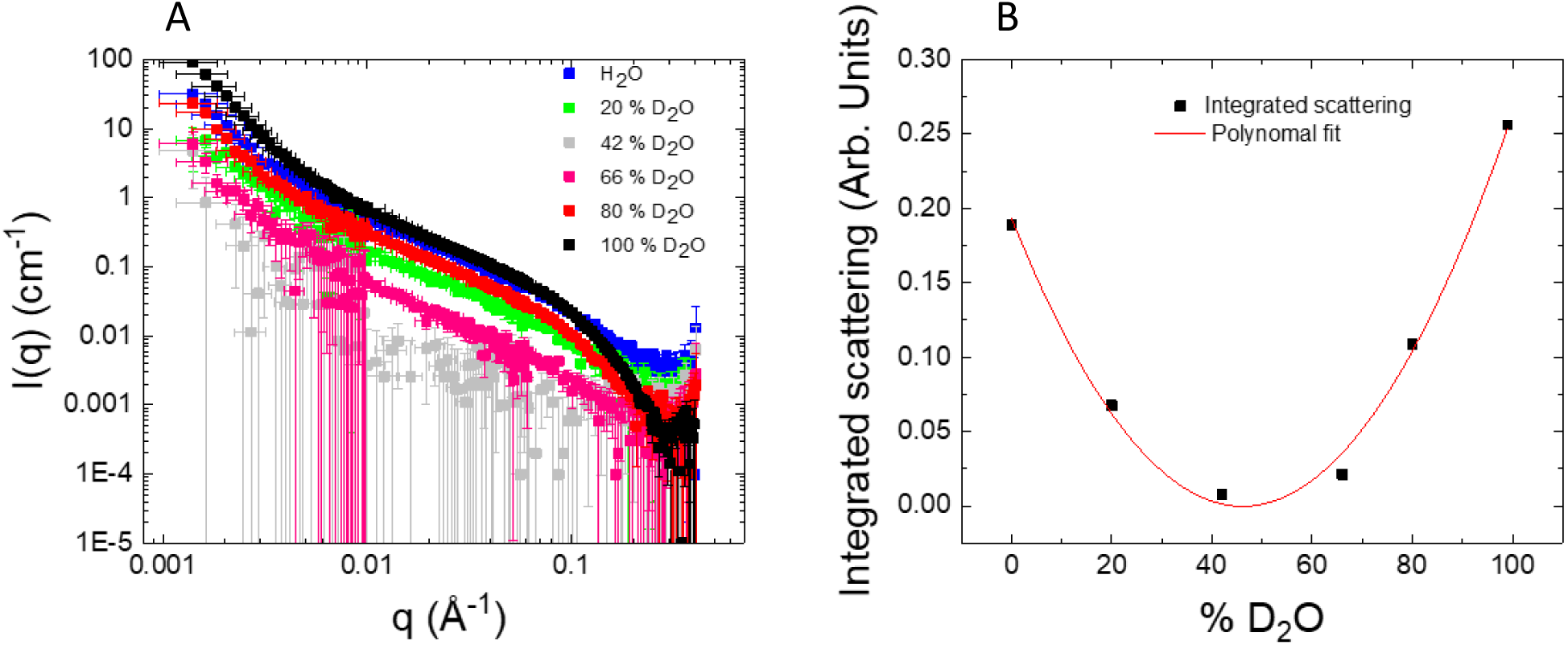
Matching out chitin in SANS experiments. **A** 10 mg/mL chitin was subjected to SANS at six different ratios of D_2_O to H_2_O, and the data were plotted as intensities (I(q)) vs. the scattering vector (q) on double-logarithmic scale. **B** The scattering for each of the six conditions was integrated, plotted and fitted with a polynomial function, to find the chitin match point at minimal scattering intensity (determined to be at 47% D_2_O).

**Fig S2.**
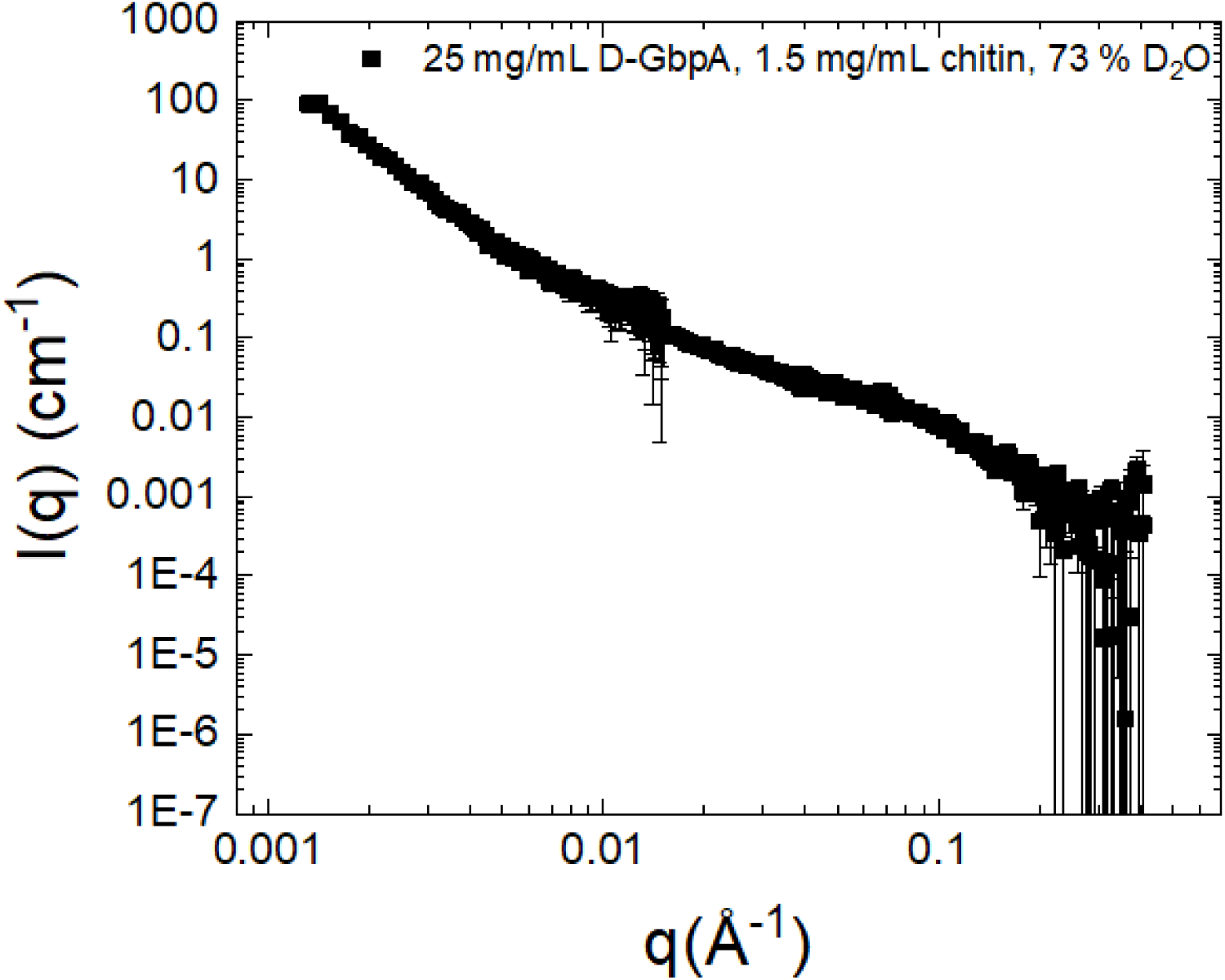
SANS data. Measurement in the middle of the chitin and D-GbpA match point yielded a slope close to the average of the slope for chitin and the complex at 47% D_2_O. Error bars represent one standard deviation.

**Fig S3.**
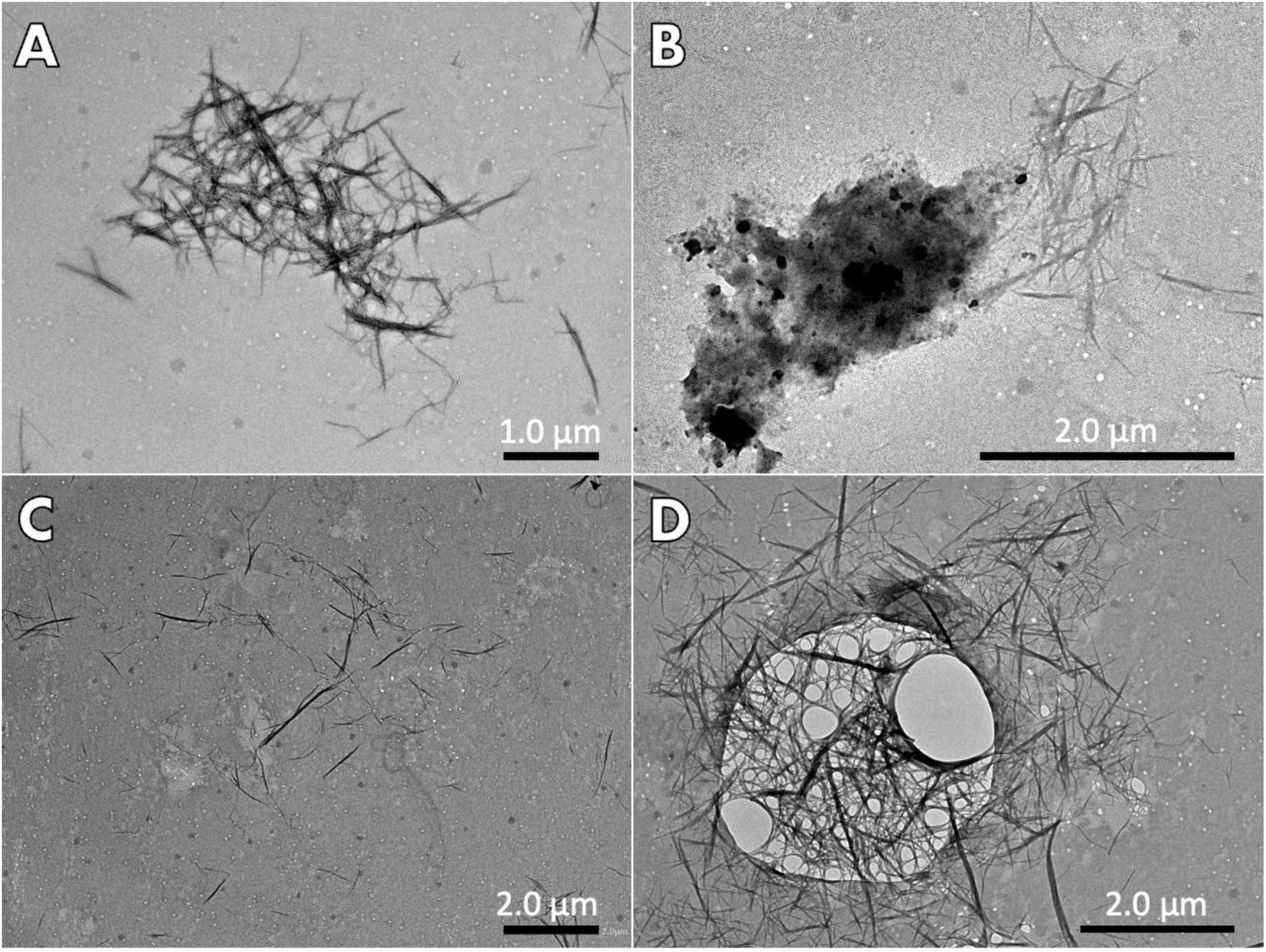
Extended TEM micrographs of chitin fibers. **A** A cluster of short chitin fibers. **B** An unknown contamination or crystallization next to a cluster of short chitin fibers. **C** Several chitin fibers spread out more evenly. **D** A cluster of chitin fibers that have been trapped in a hole in the formvar film. The concentration of chitin in these samples was 1.5 mg/mL.

**Fig S4.**
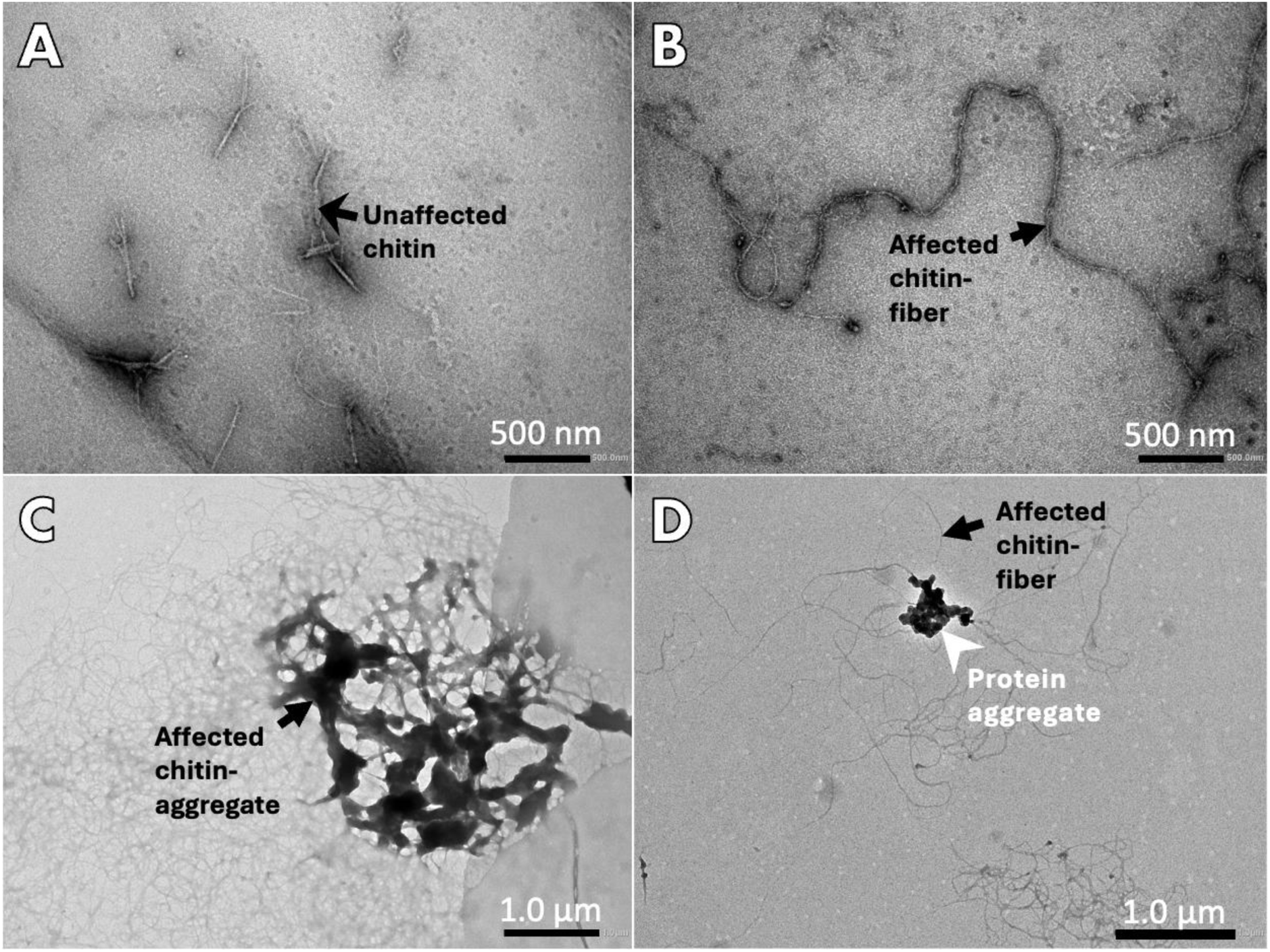
Extended TEM micrographs of chitin–GbpA complex. **A** A few relatively unaffected chitin fibers were present in this sample. **B** Close-up image of elongated fibers created by GbpA_fl_, presumably coating (and potentially linking) the chitin fibers. Note the smoother appearance compared to uncoated fibers (Fig S3). **C** Image of one of the many dense aggregates of chitin observed after complex formation with GbpA_fl_. From the cluster, a plethora of elongated fibers spread out from this denser aggregation, representing “classic” GbpA_fl_-coated chitin fibers shown in B. The hard line at the right side of this image is presumably an artifact. **D** Another GbpA-chitin cluster that is ultrastructurally characteristic of a protein aggregate, with several elongated chitin-GbpA_fl_ fibers extending from it. Additional elongated fibers are found at the bottom of the image. The concentration of chitin in these samples was 1.5 mg/mL, while the concentration of GbpA_fl_ was 400 µM.

**Fig S5.**
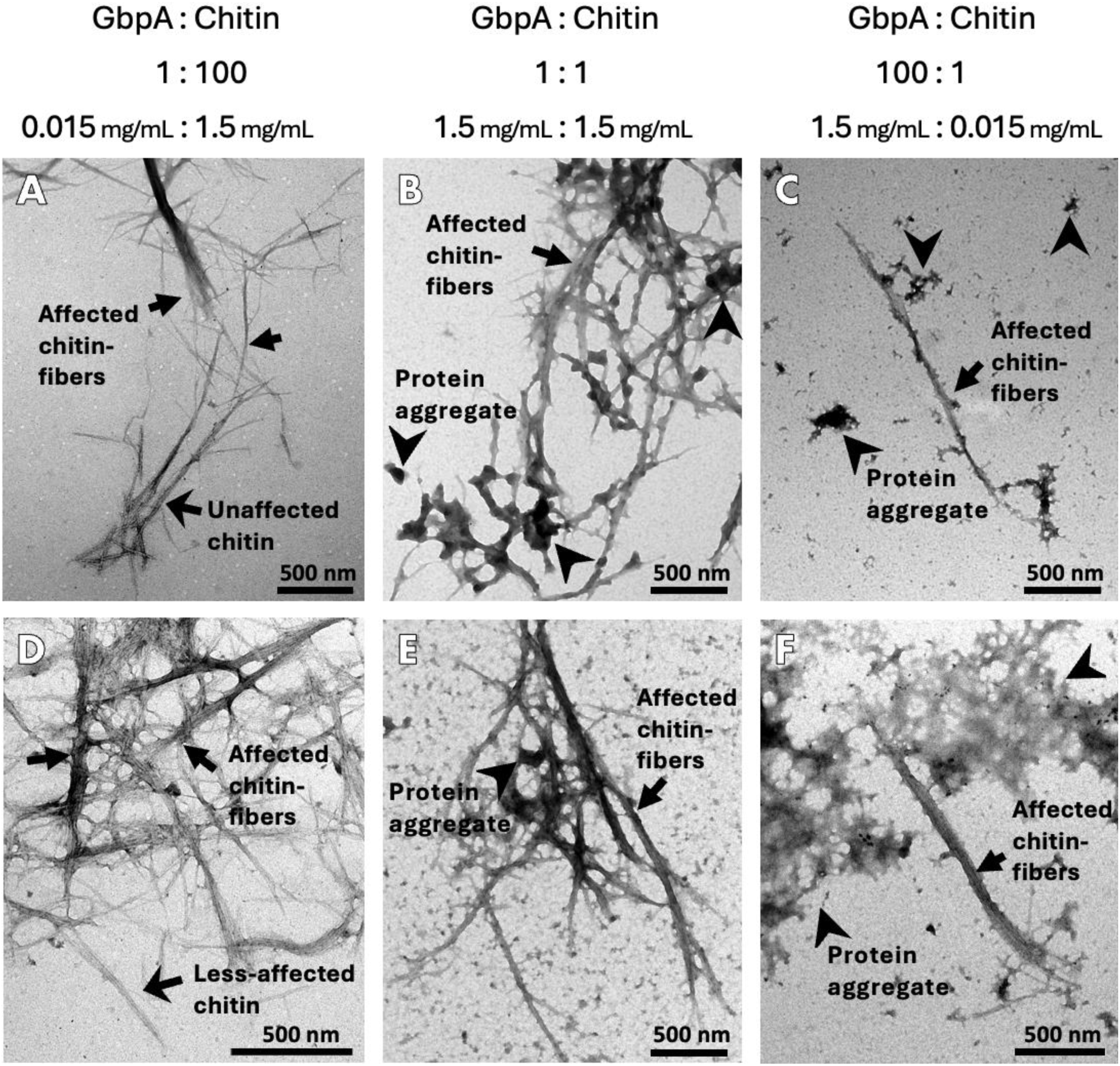
Extended TEM micrographs of different concentration ratios of chitin mixed with GbpA_fl_. (**A-C**) Shows representative results from the serial dilution experiment, while (**D-F**) are representative of results from the samples that were individually prepared, without serial dilutions. These different sample preparations had an inrteresting effect on the results. **A** In this sample containing 0.015 mg/mL GbpA: 1.5 mg/mL chitin, completely unaffected chitin fibers (broad arrow) were observed right next to GbpA_fl_-coated chitin fibers (slim arrows). Comparing this to the 1.5 mg/mL GbpA: 1.5 mg/mL chitin sample in **B**, no unaffected chitin was observed. Additionally, more numerous and denser aggregates of protein (tail-less arrows) were observed on the GbpA_fl_-coated chitin. **C** Less large clusters of chitin-fibers like were found in this sample containing 1.5 mg/mL GbpA: 0.015 mg/mL chitin (in some micrographs, one large cluster dominated the image; not shown). Most of the grid was covered in very small, dense protein aggregates that were not associated with any chitin fibers. The few chitin fibers that were found were all coated with GbpA_fl_. Essentially, this image shows a 100-fold dilution of B, with the addition of some small protein aggregates. **D** Like in A, the sample contained 0.015 mg/mL GbpA: 1.5 mg/mL chitin (however, in this case prepared directly, without prior serial dilution). No unaffected chitin fibers with sharp edges were found. **E** This sample was prepared as in B, and thus represents a replicate of B. **F** Sample comparable to C, but prepared directly, without prior serial dilution. The results are similar to C, except for the size and quantity of the protein-aggregates. The differences between **A** and **D**, but not between **B-C** and **E-F** are likely a result of the speed and affinity (with low off-rates) at which GbpA_fl_ binds to chitin. In the dilution experiment, it appears that when GbpA_fl_ and chitin were mixed in the 1:1 sample (**B**), the protein bound to the chitin so fast and with such high affinity that upon dilution, the coverage on the already-affected chitin changed very little and the added chitin remained uncoated by GbpA.

**Fig S6.**
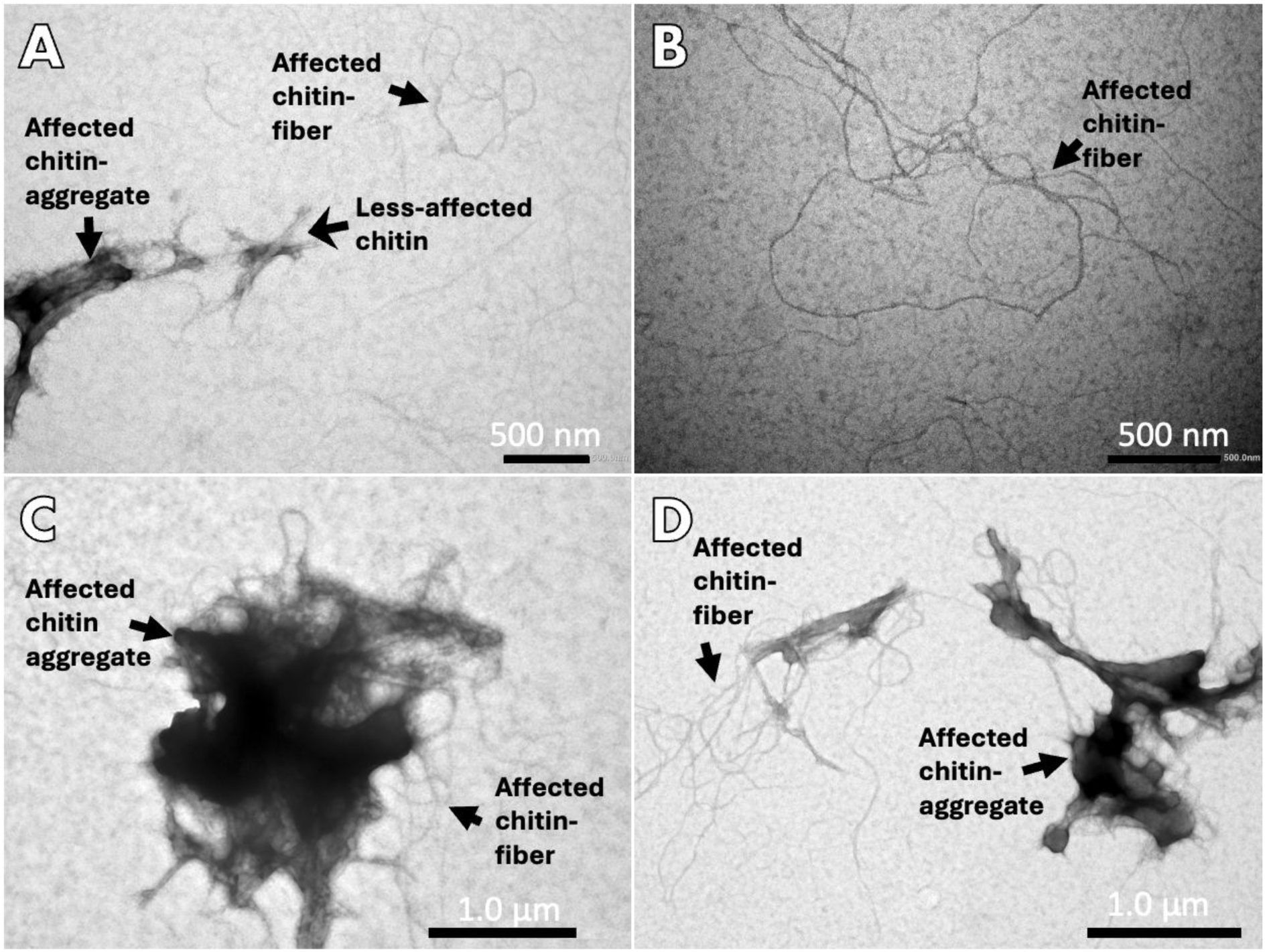
Extended TEM micrographs of chitin fibers incubated with GbpA_D1-3_. **A** A cluster of aggregated chitin affected by GbpA_D1-3_ is visible on the left, with sharper, more defined chitin fibers right next to it in the center of the image. Classic, individual, elongated looping fibers of chitin affected by GbpA_D1-3_ are also present. **B** Close-up view of individual chitin fibers coated with GbpA_D1-3_. **C** Cluster of GbpA_D1-3_-chitin aggregate with surrounding individual fibers. **D** Two smaller clusters of GbpA_D1-3_-chitin aggregates with individual fibers extending from them. The concentration of chitin in these samples was 1.5 mg/mL, while the concentration of GbpA_fl_ was 400 µM.

**Fig S7.**
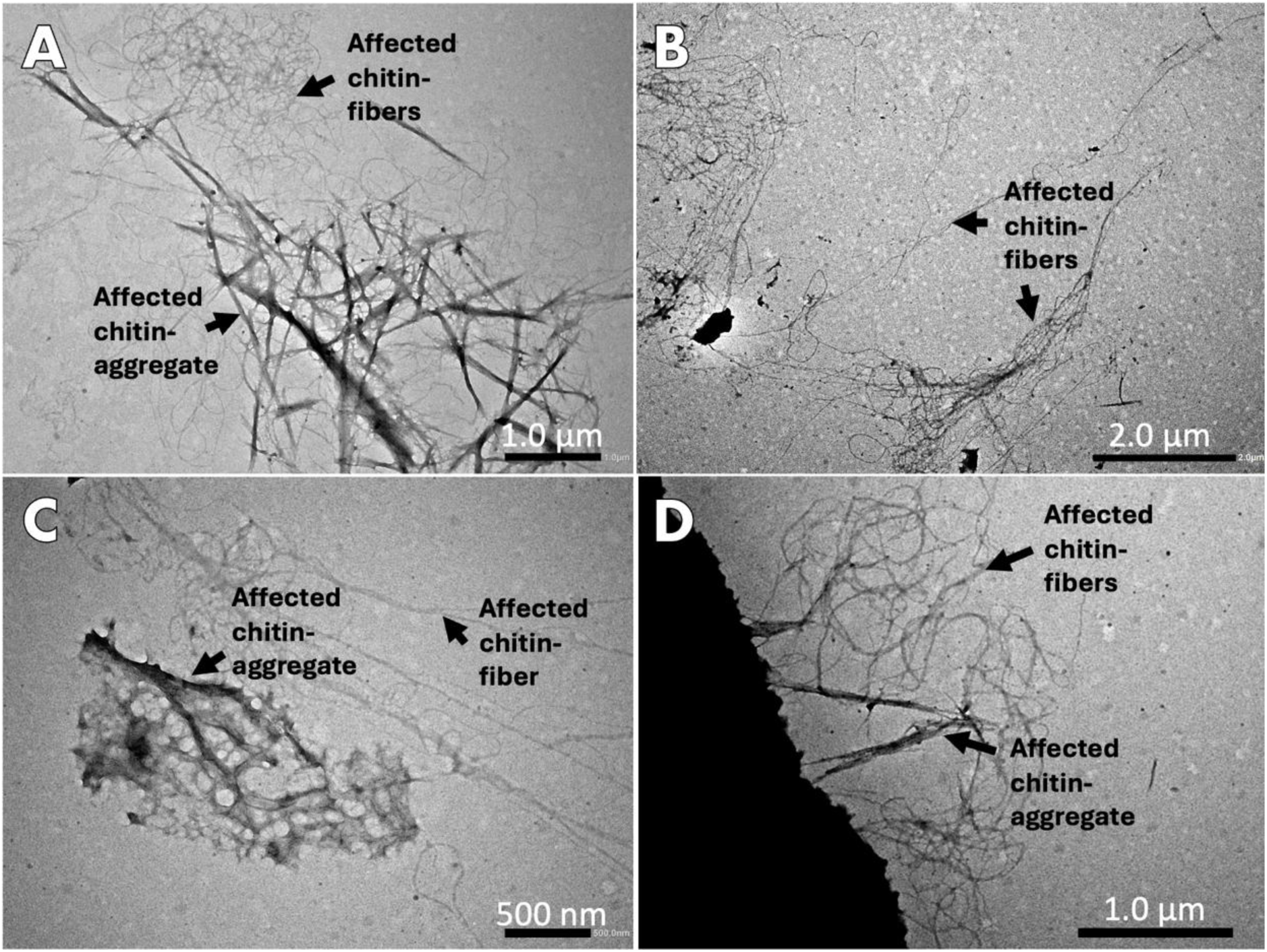
Extended TEM micrographs of chitin fibers incubated with GbpA_D1_. **A** Cluster of aggregated chitin fibers incubated with GbpA_D1_, with individual fiber loops extending above from the aggregate. Compared to chitin complexes with GbpA_fl_ (Fig S4) or GbpA_D1-3_ (Fig S5), the chitin fibers appear sharper and more defined. **B** Close-up view at individual GbpA_D1_-coated chitin fibers. **C** Cluster of GbpA_D1_-chitin aggregate with surrounding individual fibers. **D** Two smaller clusters of GbpA_D1_-chitin aggregates with individual fibers extending from them. The aggregates clearly have more jagged edges than for the longer GbpA constructs (Fig S4-5). The concentration of chitin in these samples was 1.5 mg/mL, while the concentration of GbpA_fl_ was 400 µM.

---

**Table S1:** Fitting parameters chitin cylinder model with power law.

| Sample | Length (Å) | Kuhn length (Å) | Radius (Å) | Decay exponent | $\chi^2$ | SASBDB ID |
| --- | --- | --- | --- | --- | --- | --- |
| Chitin<br>(3 mg/mL),<br>100% D<sub>2</sub>O | 5000 | 318 +/- 12 | 15.99 +/- 0.02* | 3.99 +/- 0.01 | 2.78 | SASDW42 |

*\* Note that the statistical error from the modeling is likely much lower than the actual uncertainty of the value.*

**Table S2:** Ratios of GbpA_fl_ to chitin in the dilution-series supplementary TEM experiment.

| Sample (GbpA:chitin) | 1:1000 | 1:100 | 1:10 | 1:1 | 10:1 | 100:1 | 1000:1 |
| --- | --- | --- | --- | --- | --- | --- | --- |
| GbpA<sub>fl</sub> (mg/mL) | 0.0015 | 0.015 | 0.15 | 1.5 | 1.5 | 1.5 | 1.5 |
| Chitin (mg/mL) | 1.5 | 1.5 | 1.5 | 1.5 | 0.15 | 0.015 | 0.0015 |

**Table S3:** Ratios of GbpA_fl_ to chitin in the direct-mixing supplementary TEM experiment.

| Sample (GbpA:chitin) | 1:100 | 1:1 | 100:1 |
| --- | --- | --- | --- |
| GbpA<sub>fl</sub> (mg/mL) | 0.015 | 1.5 | 1.5 |
| Chitin (mg/mL) | 1.5 | 1.5 | 0.015 |

